# Therapeutic Activity of Resolvin D1 (RvD1) in Murine MASH

**DOI:** 10.1101/2024.04.22.590633

**Authors:** Amaia Navarro-Corcuera, Yiwei Zhu, Fanglin Ma, Neha Gupta, Haley Asplund, Feifei Yuan, Scott Friedman, Brian E. Sansbury, Xin Huang, Bishuang Cai

## Abstract

**Background and Aims:** Recent studies have highlighted the beneficial effect of resolvin D1 (RvD1), a DHA-derived specialized pro-resolving mediator, on metabolic dysfunction-associated steatohepatitis (MASH), but the underlying mechanisms are not well understood. Our study aims to determine the mechanism by which RvD1 protects against MASH progression.

**Methods:** RvD1 was administered to mice with experimental MASH, followed by bulk and single-cell RNA sequencing analysis. Primary cells including bone marrow–derived macrophages (BMDMs), Kupffer cells, T cells, and primary hepatocytes were isolated to elucidate the effect of RvD1 on inflammation, cell death, and fibrosis regression genes.

**Results:** Hepatic tissue levels of RvD1 were decreased in murine and human MASH, likely due to an expansion of pro-inflammatory M1-like macrophages with diminished ability to produce RvD1. Administering RvD1 reduced inflammation, cell death, and liver fibrosis. Mechanistically, RvD1 reduced inflammation by suppressing the Stat1-Cxcl10 signaling pathway in macrophages and prevented hepatocyte death by alleviating ER stress-mediated apoptosis. Moreover, RvD1 induced *Mmp2* and decreased *Acta2* expression in hepatic stellate cells (HSCs), and promoted *Mmp9* and *Mmp12* expression in macrophages, leading to fibrosis regression in MASH.

**Conclusions:** RvD1 reduces Stat1-mediated inflammation, mitigates ER stress-induced apoptosis, and promotes MMP-mediated fibrosis regression in MASH. This study highlights the therapeutic potential of RvD1 to treat MASH.

**Impact and implications:** Metabolic dysfunction–associated steatohepatitis (MASH) is an increasing healthcare burden worldwide. Current treatments for MASH and its sequelae are very limited. Recent studies highlighted the therapeutic benefit of specialized pro-resolving mediators (SPMs), including resolvin D1 (RvD1), in liver diseases. However, the mechanisms underlying these beneficial effects are not well understood. Based on unbiased transcriptomic analyses using bulk and single-cell RNA sequencing in RvD1-treated MASH livers, we show that RvD1 suppresses Stat1-mediated inflammatory responses and ER stress-induced apoptosis, and induces gene expression related to fibrosis regression. Our study provides new mechanistic insight into the role of RvD1 in MASH and highlights its therapeutic potential to treat MASH.

**Highlights:**

- Liver RvD1 levels are decreased in MASH patients and MASH mice
- RvD1 administration suppresses Stat1-mediated inflammatory response
- RvD1 administration alleviates ER stress-induced apoptosis
- RvD1 administration induces fibrosis regression gene expression

## Introduction

Metabolic dysfunction-associated steatotic liver disease (MASLD) is one of the leading forms of chronic liver disease estimated to affect a quarter of the world’s population.^1^ Twenty to thirty percent of those with MASLD will develop metabolic dysfunction-associated steatohepatitis (MASH), a condition characterized by chronic inflammation, hepatocellular damage, and fibrosis.^1^ Without treatment, MASH can further progress to cirrhosis and hepatocellular carcinoma.^2^ Moreover, MASH is a risk factor for type 2 diabetes, obesity, and coronary artery disease.^3^ Current treatments for MASH and its sequelae are very limited. This is largely due to our lack of understanding of MASH progression, particularly the progression from steatosis to liver fibrosis, the main determinant of long-term mortality in MASH.^4^ Although the FDA has approved resmetirom, a thyroid hormone receptor β-selective agonist, as the first drug to treat MASH with fibrosis, only ∼25% of patients benefit from this treatment.^5^ Therefore, exploring new therapeutic strategies is essential.

Inflammation and cell death, two highly interconnected processes, are hallmarks of MASH and widely recognized as drivers of MASH progression.^6^ Fat overload induces lipotoxicity-mediated apoptosis in hepatocytes, leading to the release of damage-associated molecular patterns (DAMPs),^7^ which then activate resident Kupffer cells to produce inflammatory cytokines and chemokines, leading to immune cell infiltration into the liver.^8^ As MASLD progresses, monocytes are rapidly recruited to the liver where they can differentiate into macrophages and replenish Kupffer cells.^1,9^ Liver macrophages are highly plastic and dynamic in response to changes in MASH environment. By interacting with other liver cells, macrophages reshape the hepatic immune cell landscape during MASH, with direct consequences for disease severity. Excess of cytokines resulting from uncontrolled inflammation can amplify hepatocyte apoptosis.^9^ This positive feedback loop of apoptosis–inflammation leads to the activation of hepatic stellate cells (HSCs), the major collagen-producing cells in the liver, and fibrosis progression in MASH.^10^ Anti-inflammation and cell death agents such as the chemokine antagonist Cenicriviroc and the caspase inhibitor Emricasan have been tested in clinic trials of MASH, but their efficacy to reduce liver fibrosis has been limited.^11,12^ Thus, new therapeutic approaches that can actively resolve uncontrolled inflammation are needed.

Recent studies have focused on the role of inflammation resolution in tempering inflammation in chronic inflammatory diseases.^13^ Inflammation resolution is mediated in part by a class of lipid-derived autacoids called specialized pro-resolving mediators (SPMs). These include resolvins, lipoxins, maresins, and protectins, which are generated from the enzymatic metabolism of polyunsaturated fatty acids by lipoxygenases (LOX) in myeloid cells, including macrophages.^13^ SPMs act as “stop signals” of the inflammatory response to facilitate the timely resolution of inflammation while simultaneously stimulating tissue repair by blocking further immune cell infiltration and promoting efferocytosis.^13^ However, in many chronic inflammatory diseases, SPM generation is impaired, which contributes to exacerbated tissue injury.^6,14^ Indeed, the beneficial effects of SPMs have been confirmed in various inflammatory disease models and certain SPM analogues have been tested in clinical trials of infantile eczema, periodontal inflammation, and atherosclerotic cardiovascular disease, and have shown promising results.^15,16^ While the therapeutic efficacy of resolvin D1 (RvD1), a docosahexaenoic acid (DHA)–derived SPM, in reducing liver inflammation and fibrosis in chronic liver diseases,^17^ has been demonstrated, the mechanisms underlying RvD1’s beneficial effects on injured liver have not been fully characterized. In general, RvD1 elicits its pro-resolving effects via its activation of two G protein-coupled receptors—FPR2 and GPR32.^18^ RvD1 signaling through FPR2 specifically on myeloid cells has been shown to be essential for recovery from ischemic vascular injury.^19^ However, how RvD1 and its receptors are regulated in MASH is unknown.

In this study, we demonstrate for the first time that liver levels of RvD1 and FPR2 are decreased in MASH patients and MASH mouse. We further elucidate novel mechanisms by which RvD1 alleviates inflammation and cell death in MASH using bulk and single-cell RNA sequencing. Specifically, we demonstrate that RvD1 reduces liver inflammation by suppressing Stat1-mediated inflammation, mitigates ER stress-induced apoptosis, and promotes fibrosis regression. As the beneficial effects of SPM analogues have been overlooked in MASH despite being tested in clinical trials for other chronic inflammatory diseases, our study highlights the therapeutic potential of SPM analogues (*i.e.,* RvD1) in addressing MASH.

## Material and Methods

### Animal Model

Male wild-type (WT) C57BL/6J (7-to-10-week-old) mice (Jackson Laboratory, #000664) were used. The mice were put on several MASH diets as previously described: the Fibrosis and Tumors (FAT)-MASH diet (Teklad diets, TD.120528), the fructose, palmitate, cholesterol, and trans-fat diet (FPC, Teklad diets, TD.160785 PWD), the high-fat, choline-deficient, L-amino-defined diet (CDAHFD, Research diet, A06071302), and the high-fat, high-fructose, and high-cholesterol diet (FFC also called AMLN, Research diet, D17010103).^20,21^ For RvD1 administration experiment, the mice were fed the FAT-MASH diet containing 21.1% fat, 41% sucrose, and 1.25% cholesterol by weight, and a high-sugar solution (23.1 g/L d-fructose and 18.9 g/L d-glucose) for a total of 12 weeks. Simultaneously, the mice received intraperitoneal CCl_4_ (0.32 µg/g body weight) once a week. At the seven-week time point, the mice were also intraperitoneally injected with 200 mL of sterile PBS (vehicle) or RvD1 (500 ng/mouse, Cayman Chemical, #10012554) three times per week for five additional weeks while still on the FAT-MASH diet. The animals received humane care and were maintained on a 12 h light/dark cycle. All animal experiments were approved by the Animal Care and Use Committee of Icahn School of Medicine at Mount Sinai.

### Human Liver Tissues

Human liver tissue was acquired from the Liver Tissue Cell Distribution System (LTCDS) at the University of Minnesota. Phenotypic and pathological characterizations were conducted by physicians and pathologists associated with the LTCDS. The diagnostic information was included in Table S3. All human studies and analyses were performed under Icahn School of Medicine at Mount Sinai institutional review board–approved protocols.

## Results

### RvD1-FPR2 signaling is defective in human and mouse MASH livers

RvD1 is metabolized by 15-LOX from DHA (**Fig. 1A, left**). To determine whether this RvD1 biosynthesis is dysregulated in human MASH, we for the first time performed targeted LC-MS/MS to detect endogenous RvD1 levels in human livers. Strikingly, we found that liver RvD1 levels were dramatically decreased in MASH patients (**Fig. 1A, right**). Consistent with this, we found that RvD1 was reduced in the FAT-MASH mice (**Fig. 1A, right**). We then detected 17-HDHA, a 15-LOX-derived mono-hydroxy intermediate of RvD1, and found that its abundance was also decreased in both human and mouse MASH livers (**Fig. 1B**). These data indicate that the RvD1 biosynthesis pathway becomes dysregulated in MASH. Because MASH is associated with an increased ratio of M1-like macrophages to M2-like macrophages,^22^ we compared the capacity of different macrophage populations to produce RvD1. M1 and M2 polarization was induced by treating bone marrow–derived macrophages (BMDMs) with LPS + IFNγ or IL-4, respectively and was confirmed by the expression of iNOS and Arginase 1 (**Fig. 1C, left**). Interestingly, in the presence of DHA, M1-like macrophages produced less RvD1 than M2-like macrophages (**Fig. 1C, right**). Therefore, the decreased RvD1 in MASH livers may reflect the increased ratio of M1-/M2-like macrophages.

**Fig. 1.**
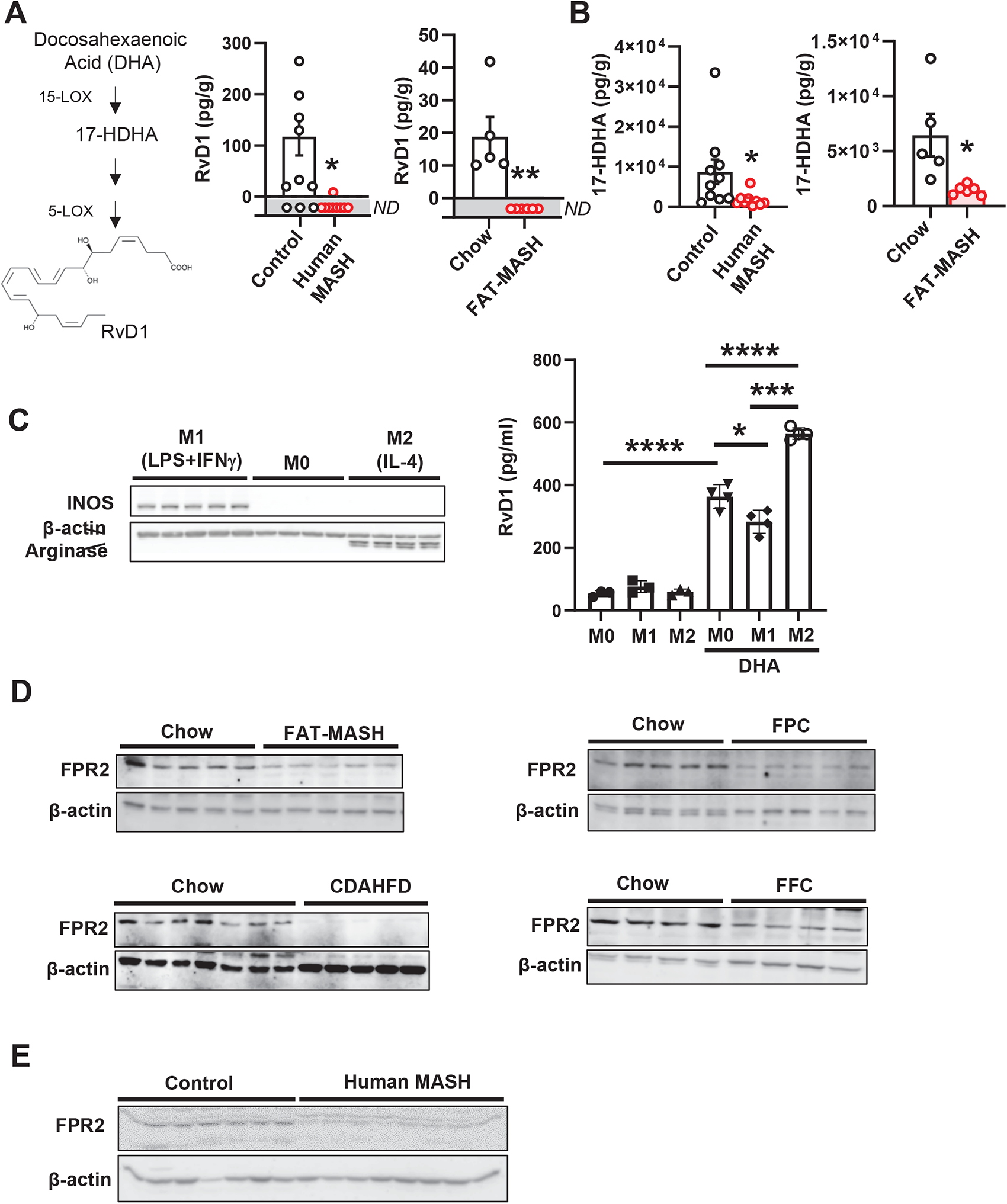
Levels of RvD1 and its receptor FPR2 are decreased in MASH livers. (A, left) Scheme showing the DHA-RvD1 conversion; (A, B) Levels of RvD1 (A) and 17-HDHA (B) in livers from control *vs.* MASH patients (n=9–10 subjects/group) and chow *vs.* FAT-MASH mice (n=5–6 mice/group) detected by LC-MS/MS (Unpaired Student’s t test \**p<*0.05, \*\**p*<0.01). (C) Immunoblots of iNOS and arginase in lysates (left panel) and ELISA analysis of RvD1 in supernatants (right panel) from polarized macrophages (n=3–4; Unpaired Student’s t test \**p*<0.05, \*\*\*\**p*<0.0001). (D, E) Immunoblot of liver FPR2 from murine MASH (D) and human MASH (E). ND, non-detected. Graph bars are presented as the mean ± SEM.

To examine whether the downstream signaling of RvD1 is affected in MASH, we determined the expression of FPR2, an RvD1 receptor, in several mouse MASH models, including the FAT-MASH, FPC-, CDAHFD-, and AMLN/FFC-induced MASH. Strikingly, we observed that liver FPR2 was significantly decreased in all the examined mouse MASH models (**Fig. 1D**). We also isolated primary hepatocytes from FAT-MASH mice to examine their FPR2 expression and found that FRP2 was reduced (**Fig. S1A**). Similar to mouse MASH livers, there was also reduced FPR2 in livers from MASH patients (**Fig. 1E**). Together, the data indicates that the RvD1-FPR2 signaling pathway becomes defective in MASH.

### RvD1 administration mitigates MASH progression

We next treated MASH mice with RvD1 to determine whether increasing RvD1 bioavailability protects against MASH. We fed WT mice with the FAT-MASH diet for 7 weeks to establish early MASH, followed by a 5-week RvD1 treatment, with continuation of the model (**Fig. 2A**). As described in previous studies with the FAT-MASH model,^20,21^ steatosis, inflammation, fibrosis, and plasma ALT were significantly induced in FAT-MASH mice compared with chow-fed mice (**Fig. 2B–D and Fig. S1B–C**). Although the liver/body weight ratio, lipid droplet area, and plasma cholesterol were not statistically different between MASH and RvD1-treated MASH mice (**Fig. S1B–D**), RvD1 administration significantly reduced plasma ALT (**Fig. 2D**), consistent with a decrease in liver injury corresponding to reduced inflammation and fibrosis, as indicated by the reduced inflammatory cells and Sirius red staining (**Fig. 2B–C**). The mRNA levels of pro-inflammatory mediators including *Tnf*, *Ccl2*, *Ifng*, and *Cxcl10* were also dramatically enhanced in MASH mice compared with chow mice, which were reversed by RvD1 (**Fig. 2E**). Moreover, RvD1-treated MASH livers expressed less *Acta2*, an HSC activation marker, and increased matrix metalloproteinase 2 (*Mmp2*), a type IV collagenase^23^ (**Fig. 2E**), in line with RvD1’s reduction of liver fibrosis.

**Fig. 2.**
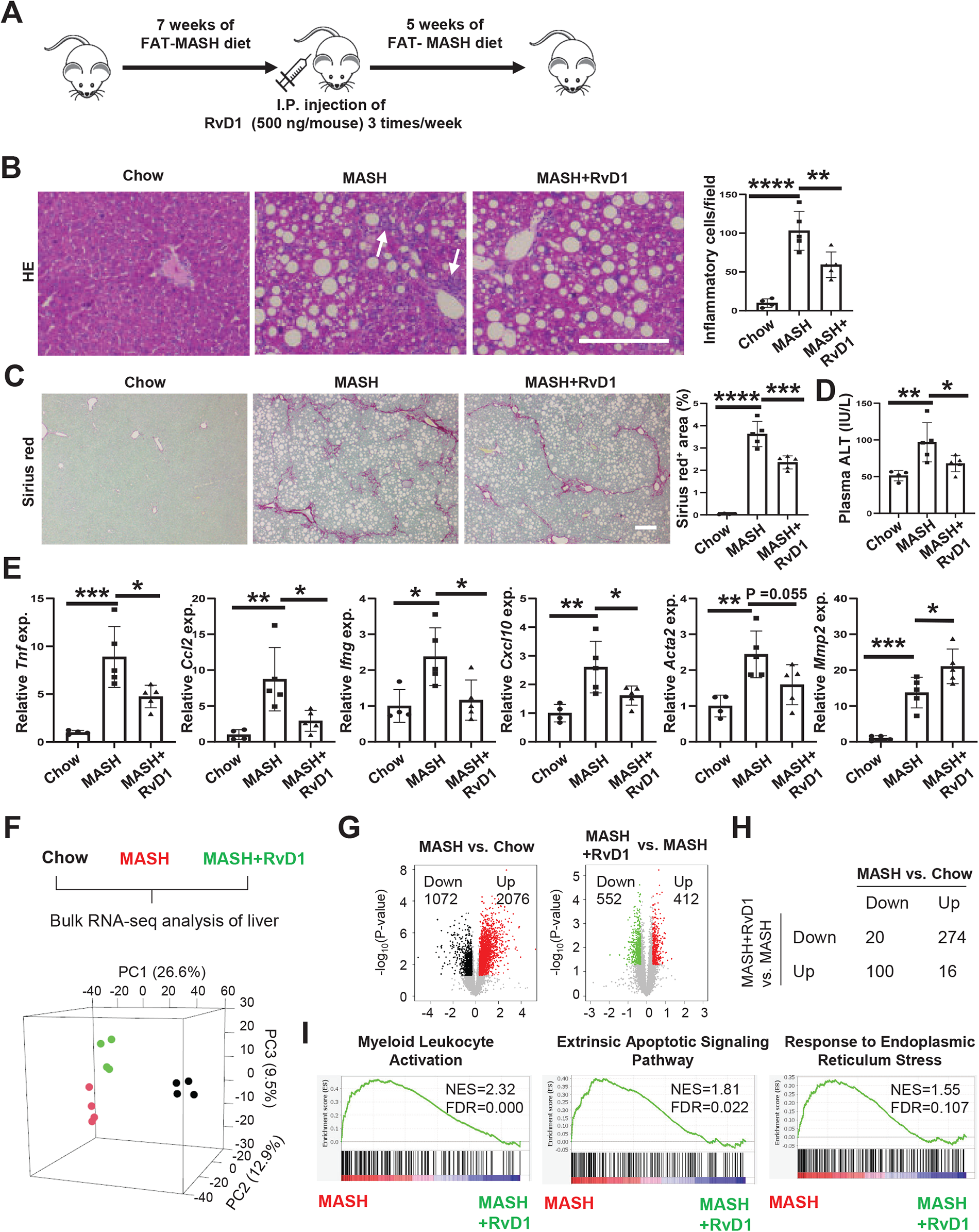
RvD1 administration attenuates MASH progression. (A) Scheme showing the experimental procedure of RvD1 administration. (B) Inflammatory cell quantification in HE-stained liver sections. Arrows indicate inflammatory cells. Bar, 200 µm. (C) Picro-Sirius Red staining in liver sections and quantification. Bar, 200 µm. (D) Plasma ALT. (E) qRT-PCR analysis for inflammatory genes in livers (In B–E, n=4–5 animals/group; one-way ANOVA with Dunnett comparison \**p*<0.05, \*\**p*<0.01, \*\*\**p*<0.001, \*\*\*\**p*<0.0001). Graph bars are presented as the mean ± SEM. (F) Illustration of bulk RNA-seq analysis (n=4 in each group) and PCA. (G) Volcano plots showing DEGs comparing MASH *vs.* chow (left) and MASH+RvD1 *vs.* MASH (right) groups. The numbers of downregulated and upregulated DESs were listed. (H) The numbers of common downregulated and upregulated genes by comparing MASH *vs.* chow and MASH+RvD1 *vs.* MASH groups were listed. (I) GSEA for the enriched gene sets that were downregulated in RvD1-treated MASH livers. The normalized enrichment score (NES) and false discovery rate (FDR) values were listed.

To further confirm the effect of RvD1 on MASH, we performed bulk RNA-seq on liver RNA isolated from chow-fed mice, and vehicle- and RvD1-treated MASH mice (**Fig. 2F**). Principal component analysis (PCA) of these RNA-seq samples revealed that the RvD1-treated MASH livers (green dots) were separated from the MASH livers (red dots), and more importantly, they were closer to the chow/healthy livers (black dots) compared with MASH livers (**Fig. 2F**), indicating that RvD1 alleviates MASH. Next, we obtained the differentially expressed genes (DEGs, by P-value<0.05 and fold-change>1.25 based on FPKM+1 values) between different groups (**Fig. 2G**) and found that DEGs upregulated in the MASH group compared to the chow group were mostly (n=274) downregulated in MASH+RvD1 as compared to the MASH group (**Fig. 2H**). Similarly, DEGs downregulated in the MASH group compared to the chow group were mostly (n=100) upregulated in the MASH+RvD1 group compared to the MASH group (**Fig. 2H**). These results indicate that RvD1 treatment inhibits MASH progression. Comparison by gene set enrichment analysis (GSEA) between MASH+RvD1 and MASH groups revealed that myeloid leukocyte activation, extrinsic apoptotic signaling pathway, and response to endoplasmic reticulum stress were downregulated in RvD1-treated MASH livers (**Fig. 2I**). Altogether, these results indicate that RvD1 protects against MASH.

### RvD1 eliminates inflammation by suppressing Stat1-Cxcl10 signaling in macrophages

We next explored the mechanism by which RvD1 suppresses inflammation in MASH. The RNA-seq data showed that *Cxcl10,* an IFNγ-inducible chemokine,^24^ was the top RvD1-downregulated inflammatory gene (**Figs. 3A and S1E**), indicating that RvD1 may reduce inflammation via suppressing *Cxcl10* expression in MASH. *Cxcl10* expression can be induced by the transcription factor Stat1, the major Stat protein activated by IFNγ.^25^ Consistent with this concept, Stat1 activation, as determined by phosphorylated Stat1 (p-Stat1), was decreased in RvD1-treated MASH livers (**Fig. 3B**). As Cxcl10 is highly expressed in liver mononuclear phagocytes, including macrophages (**Fig. S1F**),^26^ we examined the effect of RvD1 on Stat1 activation in liver macrophages and found that p-Stat1 was decreased in Mac2^+^ macrophages in RvD1-treated MASH livers (**Fig. 3C**). The decreased Stat1 activation and Cxcl10 expression are likely due to decreased IFNγ expression in RvD1-treated MASH mice as indicated in our RNA-seq (**Fig. 3A**). Since activated T cells are the major cell type producing IFNγ, and SPMs, including RvD1 reduce IFNγ production in human circulating T cells,^27^ we investigated whether RvD1 suppresses IFNγ production in liver T cells. We isolated primary liver T cells from WT mice and activated these T cells with anti-CD3/CD28 antibodies^28^ to induce IFNγ production in the absence or presence of RvD1. As expected, we found that IFNγ production was dramatically induced in activated T cells and, similar to its effect on human circulating T cells,^27^ RvD1 suppressed IFNγ production in mouse liver T cells (**Fig. 3D**). We next determined the effect of IFNγ in macrophages and found that IFNγ activated Stat1 and induced Cxcl10 in liver resident Kupffer cells (**Fig. 3E**). MASH can induce bone marrow–derived monocyte infiltration and differentiation in livers (infiltrated macrophages),^29^ and similar to Kupffer cells, the Stat1-Cxcl10 pathway was also upregulated by IFNγ in BMDMs (**Fig. 3F**). These data indicate that RvD1 modulates the cellular crosstalk between T cells and macrophages to suppress IFNγ/Cxcl10-mediated inflammation.

**Fig. 3.**
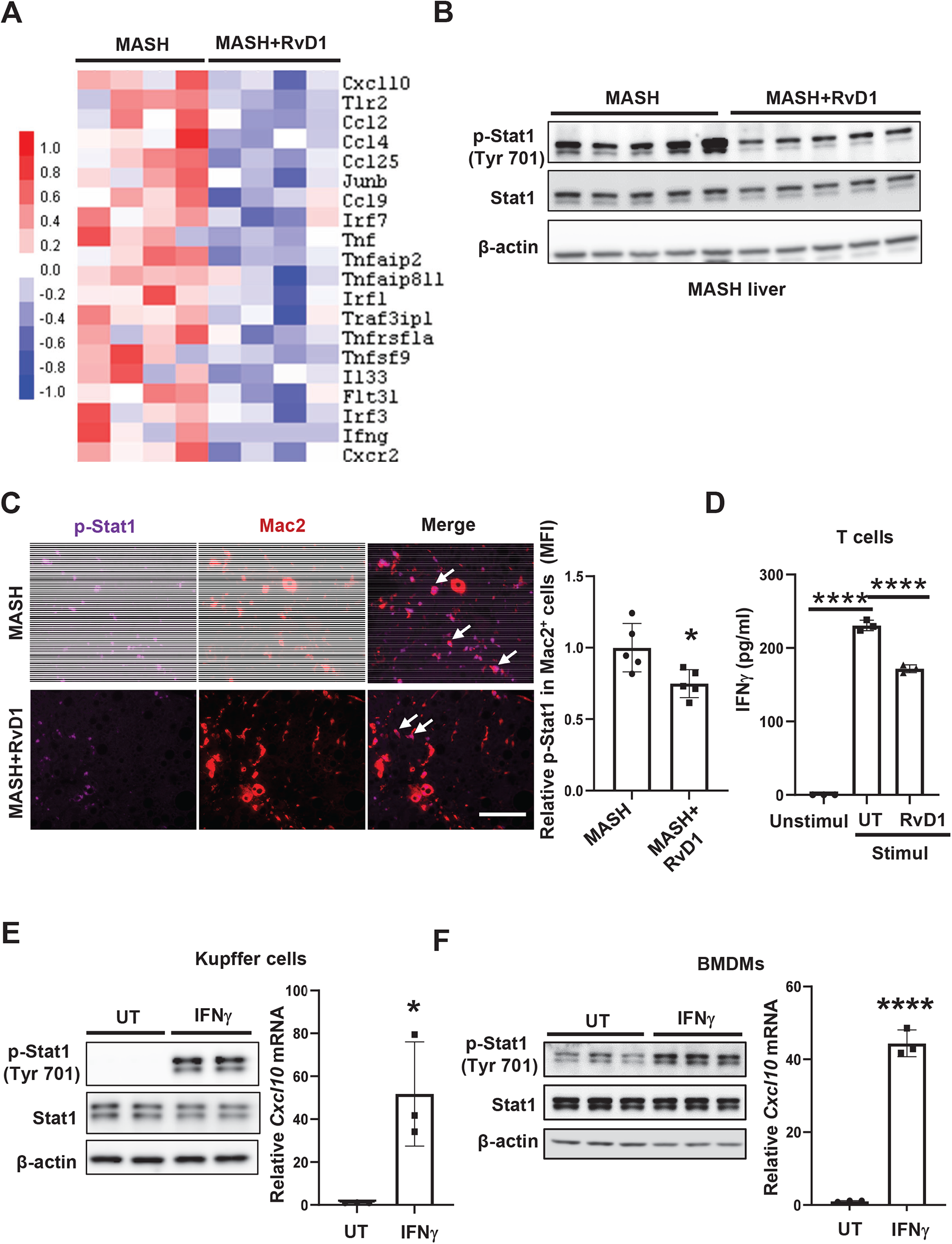
RvD1 attenuates inflammation by inhibiting Stat1 activation. (A) Heatmap showing the expression of inflammatory genes. A color scale of Z-score was shown on the left. (B) Immunoblots of phospho-Stat1 (p-Stat1) and Stat1 in FAT-MASH livers. (C) Co-immunostaining showing p-Stat1 levels in Mac2^+^ macrophages in FAT-MASH livers. The mean fluorescence intensity (MFI) of p-Stat1 was quantified by Image J (n=5 mice/group; Unpaired Student’s t test \**p*<0.05). White arrows indicate the co-staining of p-Stat1 and Mac2. Bar, 100 µm. (D) Primary T cells were isolated from WT mice and were unstimulated (Unstimul) or stimulated with anti-CD3/CD28 antibodies (Stimul) for 18 h in the absence (untreated, UT) or presence of 50 nM RvD1. IFNγ in supernatants was measured by ELISA (n=3 mice/groups; one-way ANOVA with Dunnett comparison \*\*\*\**p*<0.0001). (E, F) Immunoblots of p-Stat1 and Stat1 (left panel) and qRT-PCR for *Cxcl10* (right panel) in Kupffer cells (E) and BMDMs (F), treated with IFNγ or untreated (UT) (n=3 mice/group; Unpaired Student’s t test \**p<*0.05, \*\*\*\**p*<0.0001). Graph bars are presented as the mean ± SEM.

### RvD1 reduces cell death by suppressing the ROS-CHOP pathway in hepatocytes

In addition to inflammation, RNA-seq from RvD1-treated MASH livers demonstrated that RvD1 also suppresses pathways underlying ER stress and cell death (**Fig. 2I**). We therefore explored the role of RvD1 in cell death in MASH. RvD1 suppressed the expression of cell death–related genes including *Zbp1*, *Gsdmd*, and *Bax* in MASH livers (**Fig. 4A**). Levels of cleaved caspase-3 and TUNEL, markers of cell death in MASH livers, were attenuated in RvD1-treated MASH livers (**Fig. 4B–C**). Since hepatocyte apoptosis strongly correlates with MASH progression,^30^ we next determined whether RvD1 prevents apoptosis in primary hepatocytes. Primary hepatocytes were incubated with palmitic acid,^31^ a MASH-relevant saturated fatty acid that can elicit lipotoxicity and apoptotic cell death, in the absence or presence of RvD1. As expected, palmitic acid induced apoptosis, as indicated by the increased cleaved caspase-3; however, RvD1 prevented palmitic acid–induced cell death in primary hepatocytes (**Fig. 4D**).

**Fig. 4.**
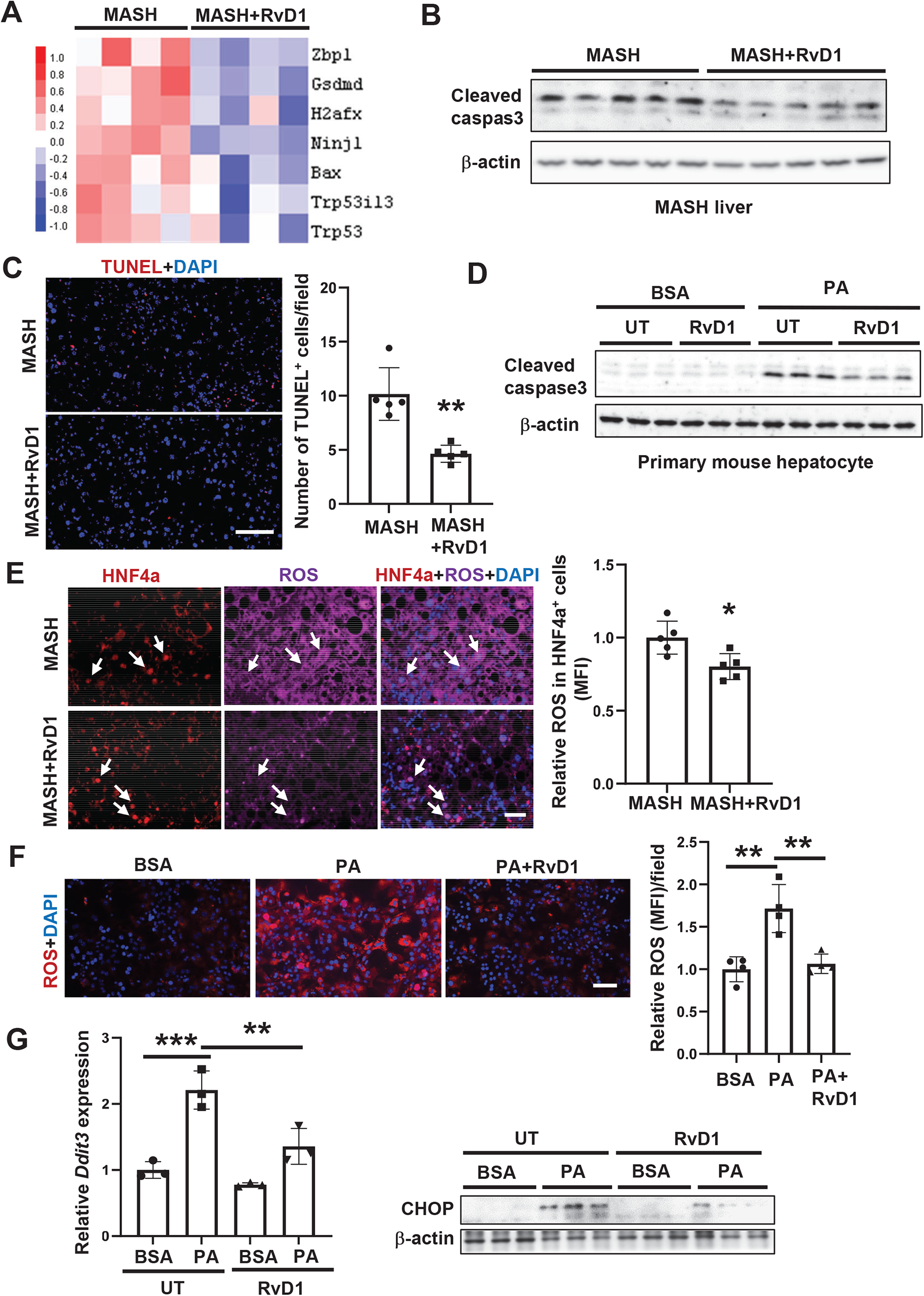
RvD1 inhibits ROS production, ER stress, and apoptosis. (A) Heatmap depicting the expression of cell death–related genes. (B) Immunoblot of cleaved caspase-3 in FAT-MASH livers. (C) TUNEL staining in FAT-MASH livers. The number of TUNEL^+^ cells was quantified by Image J (n=5 mice/group; Unpaired Student’s t test \*\**p*<0.01). Bar, 100 µm. (D) Immunoblot of cleaved caspase-3 in primary mouse hepatocytes treated with BSA or 100 µM palmitic acid (PA) ± 50 nM RvD1 for 24 h. (E) Co-immunostaining showing ROS levels in HNF4α^+^ cells in MASH livers. The MFI of ROS was quantified by Image J (n=5 mice/group; Unpaired Student’s t test \**p*<0.05 compared with FAT-MASH group). White arrows indicate the co-staining of ROS and HNF4α. Bar, 50 µm. (F) ROS staining in primary mouse hepatocytes treated with BSA or PA ± RvD1. The MFI of ROS was quantified by Image J (n=4; one-way ANOVA with Dunnett comparison \*\**p*<0.01). Bar, 50 µm. (G) qRT-PCR (left panel) and immunoblot (right panel) of CHOP in primary mouse hepatocytes treated with BSA or PA ± RvD1 (n=3, Unpaired Student’s t test \**p*<0.05, \*\**p*<0.01). Graph bars are presented as the mean ± SEM.

To determine whether the effect of RvD1 on cell death is specific to palmitic acid–mediated lipotoxicity, primary hepatocytes were exposed to TNFα/Jo2, to induce death receptor–mediated apoptosis.^32^ Interestingly, RvD1 suppressed death receptor–induced cell death as well (**Fig. S1G**). We next sought to determine the mechanism by which RvD1 suppresses cell death. Given that RvD1 has been shown to reduce reactive oxygen species (ROS) production^33^ and that ROS can promote ER stress-induced apoptosis,^34,35^ we assayed ROS levels in HNF4α^+^ hepatocytes from MASH livers and found that ROS was significantly decreased in hepatocytes from RvD1-treated MASH livers (**Fig. 4E**). Consistent with the *in vivo* observation, RvD1 inhibited palmitic acid–induced ROS production in primary hepatocytes (**Fig. 4F**). To link RvD1-reduced ROS to ER stress, we examined whether RvD1 regulates palmitic acid–induced ER stress in primary hepatocytes and found that palmitic acid induced the expression of C/EBP homologous protein (CHOP, encoded by *Ddit3*), an ER stress marker, however, RvD1 blocked this induction (**Fig. 4G**). This data suggests that RvD1 suppresses cell death by alleviating ROS/CHOP-mediated apoptosis.

### RvD1 suppresses the activation of immune cells in MASH

To further explore the regulation of RvD1 in immune cell populations, we carried out single-cell RNA-seq in non-parenchymal cells isolated from MASH livers. A total of 28,059 single-cell transcriptomes were analyzed (chow: 3,017; MASH: 11,599; MASH+RvD1: 13,443). Clustering visualized on uniform manifold approximation and projection (UMAP) revealed eight major clusters of cells (**Figs. 5A and S2A**), including endothelial cells (11,503), dendritic cells (1,267), macrophages (10,828), hepatic stellate cells (HSCs, 2,326), T cells (472), B cells (171), hepatocytes (692), and cholangiocytes (800), and was annotated with lineage-specific marker genes (**Fig. 5B**).

**Fig. 5.**
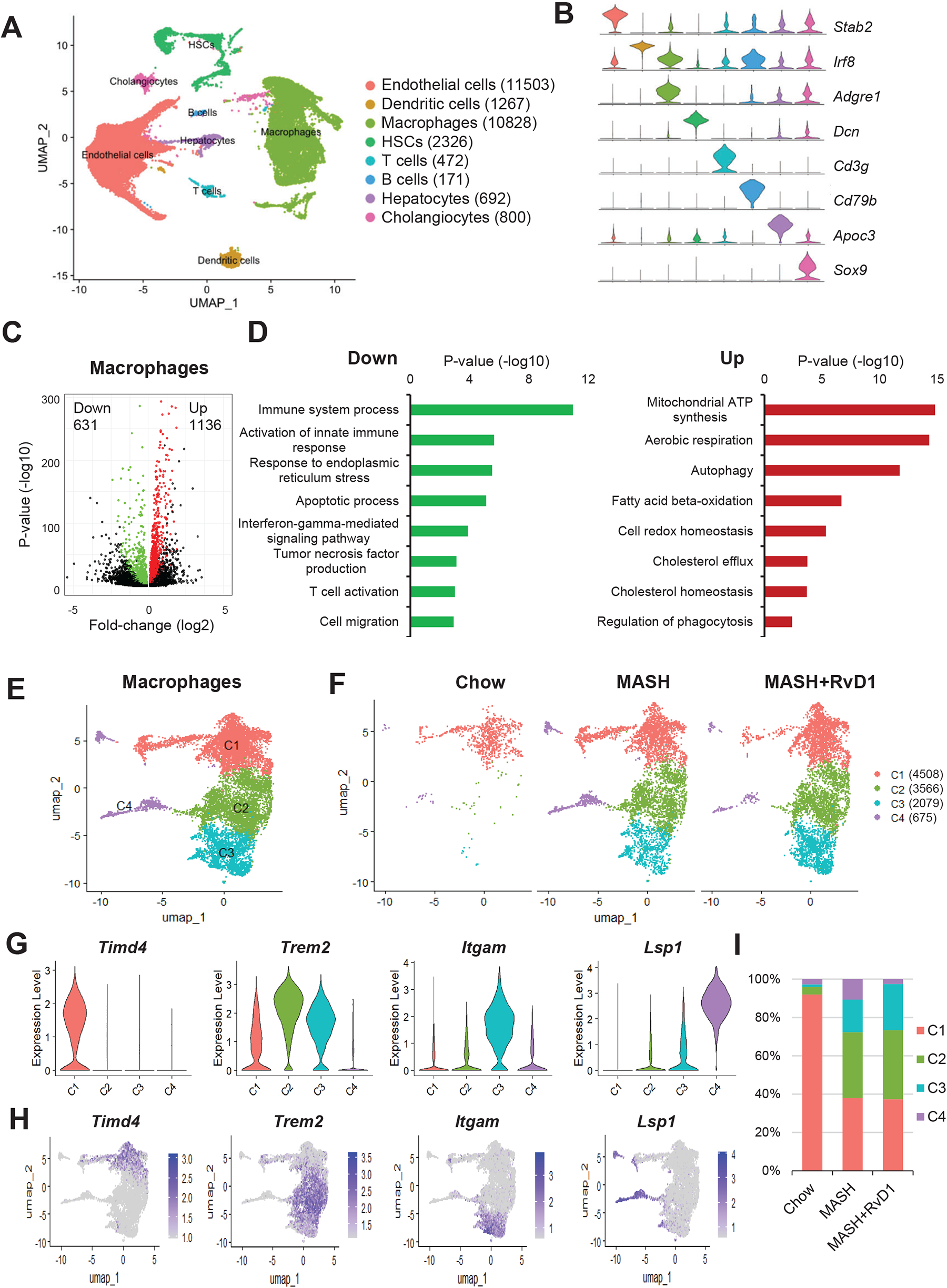
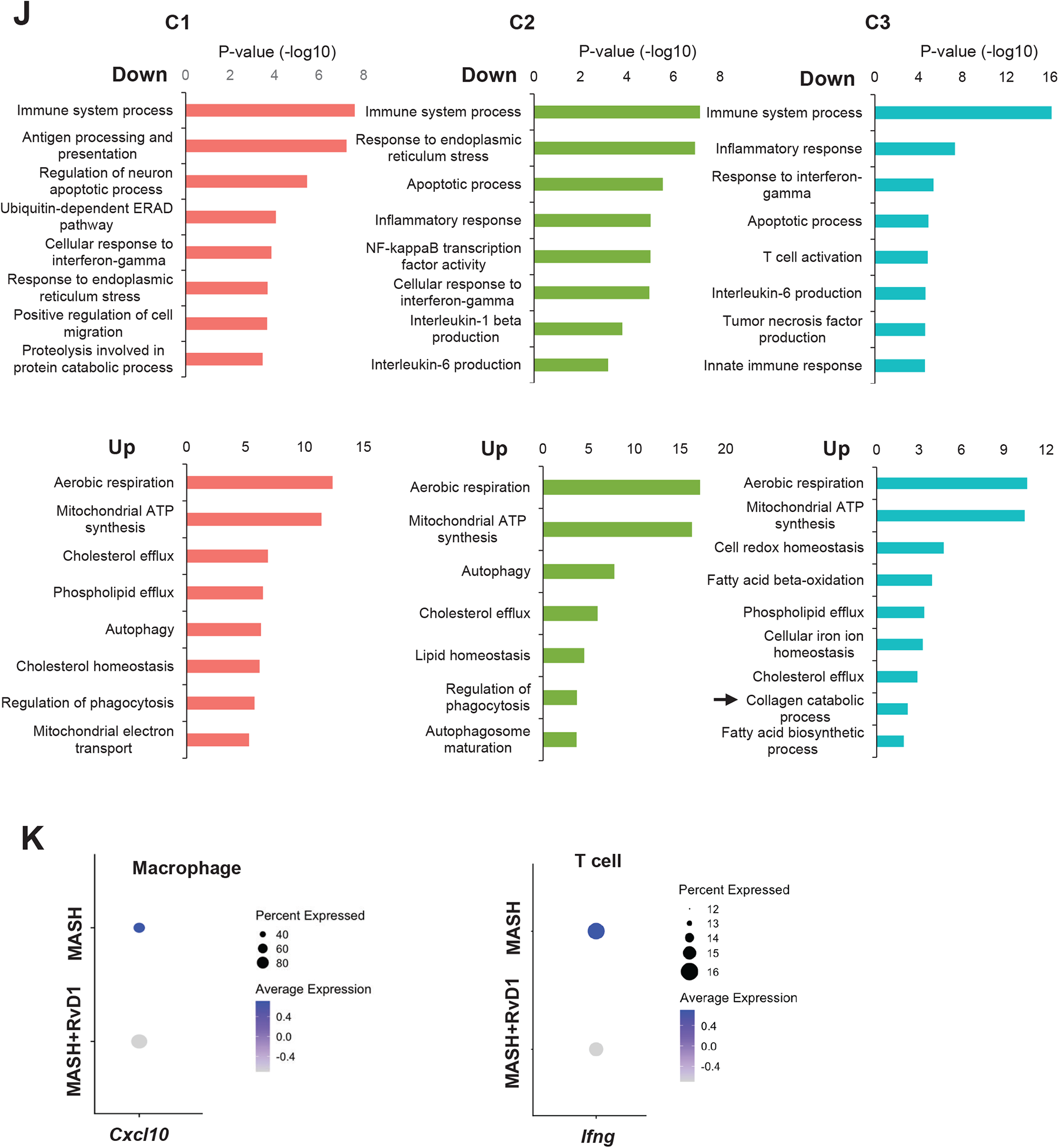
Single-cell RNA-seq analysis reveals that RvD1 suppresses macrophage activation in MASH liver. (A) UMAP visualization of liver cells based on 28,059 single-cell transcriptomes. Cell counts for each annotated cell type were indicated in parentheses. (B) Violin plots showing representative marker gene expression for each cell type. (C) Volcano plot showing macrophage DEGs by comparing MASH+RvD1 *vs.* MASH groups. (D) Gene ontology (GO) analysis for the downregulated (left) and upregulated (right) DEGs in macrophages. (E, F) UMAP plots for all the macrophages (E) and C1–C4 macrophage subclusters (F) in chow, FAT-MASH, and FAT-MASH+RvD1 groups. (G, H) Violin plots (G) and feature plots (H) for the marker gene expression in each macrophage subcluster. (I) Percentage of cell counts in each macrophage subcluster grouped by different mouse groups. (J) GO analysis for the downregulated and upregulated DEGs in C1–C3 macrophage subclusters by comparing MASH+RvD1 *vs.* MASH groups. (K) Dot plots showing *Cxcl10* (left) and *Ifng* (right) expression in macrophages and T cells, respectively.

Since macrophages are the most prominent immune cells in MASH livers, we determined the effect of RvD1 on macrophages. We performed DEG analysis in macrophages and identified 631 downregulated and 1,136 upregulated genes (P-value<0.05, percentage of expression >25% in both groups) by comparing MASH+RvD1 *vs.* MASH (**Fig. 5C**). Gene ontology (GO) analysis for these DEGs demonstrated that biological processes including immune system process, response to ER stress, IFNγ−mediated signaling pathway, and TNF production were significantly enriched in downregulated DEGs (**Fig. 5D, left**). In contrast, mitochondrial ATP synthesis, aerobic respiration, fatty acid beta-oxidation, and cholesterol efflux were significantly enriched in upregulated DEGs (**Fig. 5D, right**). Macrophages are highly heterogeneous in MASH livers; therefore, we subclustered macrophages into four populations by comparing the marker genes including *Timd4*, *Trem2*, *Itgam* (CD11b), and *Lsp1* (**Fig. 5E**). As predicted, chow mice predominantly harbored Timd4^high^ resident macrophages (C1), while both MASH and RvD1-treated MASH mice had infiltrated Trem2^high^ (C2), CD11b^high^ (C3), and Lsp1^high^ (C4) macrophages (**Fig. 5F–H**). The C3 population was increased in RvD1-treated MASH mice compared with MASH (**Fig. 5I**). By GO analysis for the DEGs in the individual macrophage subclusters (C1–C3; C4 was not included due to limited cell numbers), there was downregulation of immune system process, cellular response to IFNγ, and apoptotic process, while mitochondrial function and cholesterol efflux were upregulated in these three macrophage subclusters in RvD1-treated MASH livers (**Fig. 5J**). Of note, the collagen catabolic process was upregulated in CD11b^high^ C3 macrophages, indicating that C3 macrophages may protect against fibrosis (**Fig. 5J, arrow labeled**). The enriched genes involved in this collagen catabolic process included *Adam15*, *Ctsl*, *Mmp12*, and *Mmp14* (**Fig. S2B**). Intriguingly, C3 macrophages also highly express osteopontin (Spp1) (**Fig. S2C**), which was recently shown to protect against MASH progression.^36^ Spp1^high^ macrophages express higher levels of genes regulating extracellular matrix remodeling, such as MMPs,^36,37^ indicating the potential role of Spp1 in facilitating the resolution of liver fibrosis. Similar to the effect in macrophage, RvD1 treatment downregulated pathways involved in TNF production, antigen processing and presentation, and apoptosis, while upregulating pathways involved in mitochondrial ATP synthesis and aerobic respiration in dendritic cells, a myeloid lineage closely related to macrophages (**Fig. S2D–E**). Consistent with our bulk RNA-seq showing reduced *Cxcl10* and *Ifng* expression in RvD1-treated MASH livers (**Fig. 3**), *Cxcl10* expression in liver macrophages and *Ifng* expression in T cells were lower in RvD1-treated MASH mice (**Fig. 5K**). These results confirm that RvD1 eliminates inflammation by suppressing the activation of immune cells in MASH.

### RvD1 promotes *Mmp* expression and suppresses HSC activation in MASH

Although macrophage-derived proinflammatory mediators (*i.e.*, Cxcl10 and TNF) and damage-associated molecular patterns (DAMPs) released from dying hepatocytes have been recognized as important features that promote fibrosis progression in chronic liver disease,^38^ we sought to explore whether RvD1 plays a role in fibrosis regression in addition to its role in reducing inflammation and cell death. MMPs are well known for their anti-fibrotic effect due to their ability to degrade extracellular matrix (ECM) proteins and accelerate fibrosis regression.^39^ We then analyzed the expression profile for the major MMPs that have been shown to regulate liver fibrosis, including *Mmp2*, *Mmp9*, and *Mmp12*,^23,40,41^ by single-cell RNA-seq of MASH livers. *Mmp2* was predominantly expressed in HSCs, while *Mmp9* and *Mmp12* were specifically expressed in macrophages, particularly in C3 macrophages (**Fig. 6A and S2B**).

**Fig. 6.**
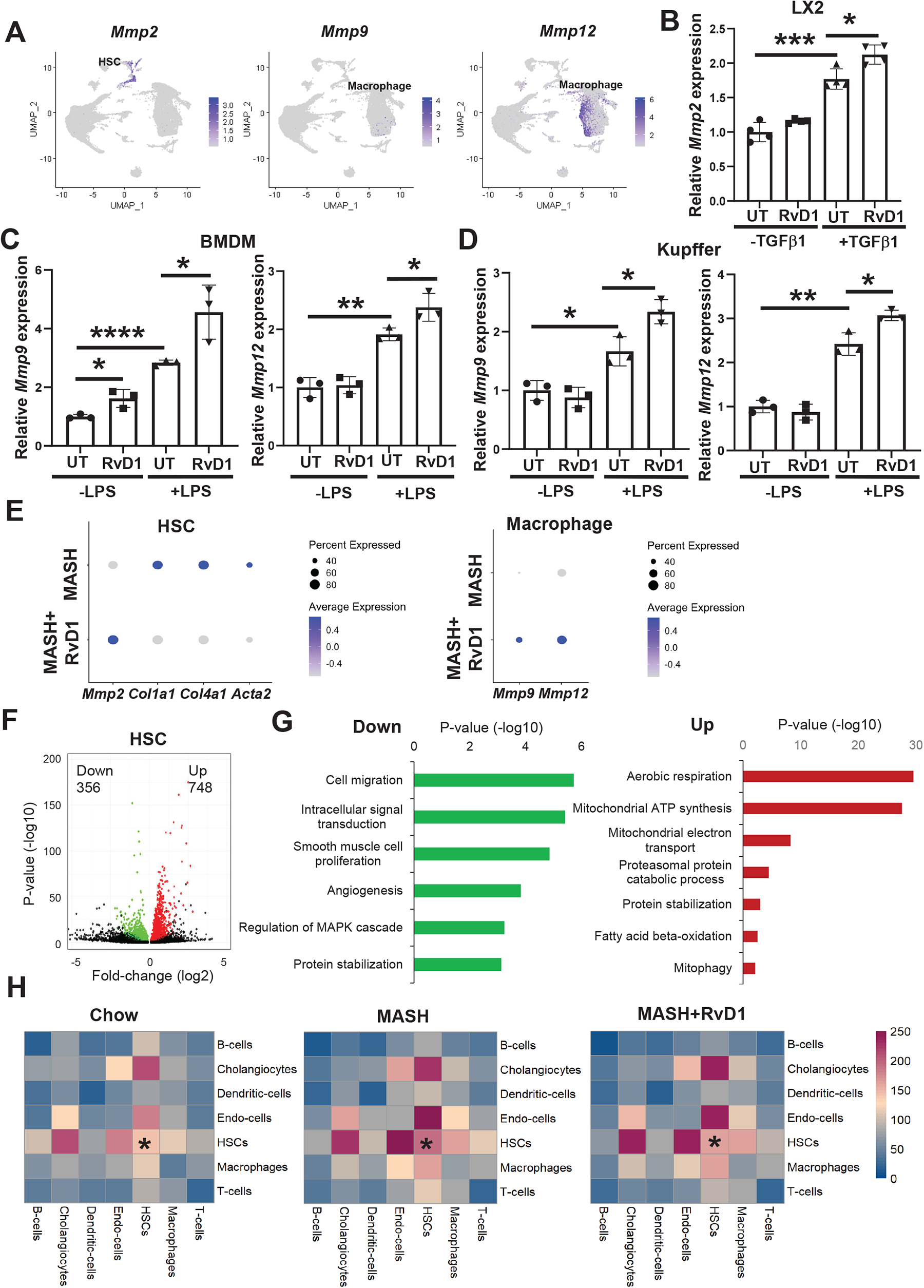
RvD1 promotes *Mmp* expression and suppresses HSC activation in MASH. (A) Feature plots showing expression of *Mmp2*, *Mmp9*, and *Mmp12* genes. (B) LX2 cells were activated with 20 ng/mL TGFβ1 ± 50 nm RvD1 for 18 h. *Mmp2* mRNA levels were detected by qRT-PCR (n=4; Unpaired Student’s t test \**p*<0.05, \*\*\**p*<0.001). (C, D) BMDMs (C) or Kupffer cells (D) were activated with 50 ng/mL LPS ± 50 nM RvD1 for 18 h. *Mmp9* and *Mmp12* mRNA levels were detected by qRT-PCR (n=3; Unpaired Student’s t test \**p*<0.05, \*\**p*<0.01, \*\*\*\**p*<0.0001). (E) Dot plots showing selected genes in HSCs and macrophages. (F) Volcano plot showing DEGs in HSCs by comparing MASH+RvD1 *vs.* MASH groups. (G) GO analysis for the downregulated (left) and upregulated (right) DEGs in HSCs. (H) Cell-cell interaction heatmap between different cell types from CellphoneDB analysis. The asterisk indicates the self-interaction of HSCs. A color scale of interaction counts was shown on the right. Graph bars in B–D are presented as the mean ± SEM.

We next determined whether RvD1 directly regulates *Mmp* expression. We found that RvD1 induced *Mmp2* expression in TGFβ1-activated LX2 cells, a human HSC line (**Fig. 6B**). Similarly, RvD1 enhanced both *Mmp9* and *Mmp12* expression in LPS-activated BMDMs and Kupffer cells (**Fig. 6C–D**). We then determined whether RvD1 modulates *Mmp* expression in MASH livers. In line with our *in vitro* studies, RvD1 promoted *Mmp2* expression in HSCs from MASH livers (**Fig. 6E**). Interestingly, the increased *Mmp2* was associated with decreased expression of collagen genes (*i.e.*, *Col1a1*, *Col4a1)* as well as *Acta2* (**Fig. 6E**). Similar to HSCs, macrophages had increased expression of *Mmp9* and *Mmp12* in RvD1-treated MASH livers (**Fig. 6E**). The increased *Mmp* indicates that RvD1 may promote fibrosis regression. To determine if RvD1 has a broad effect on HSCs, we next performed DEG analysis in HSCs and identified 356 downregulated and 748 upregulated genes by comparing RvD1-treated MASH vs. MASH (**Fig. 6F**). GO analysis for these DEGs revealed that cell migration and proliferation were downregulated, while aerobic respiration and mitochondrial ATP synthesis were upregulated in HSCs from RvD1-treated MASH livers (**Fig. 6G**). Moreover, consistent with a recent study showing that the prominent HSC autocrine signaling circuit is a key driver for MASH fibrosis,^21^ our CellphoneDB analysis successfully captured an increased HSC-HSC interaction in the FAT-MASH mice compared with chow mice, and interestingly, we demonstrated that RvD1 alleviated HSC-HSC contacts in MASH (**Fig. 6F**). Together, our data suggests that RvD1 protects against liver fibrosis by inducing MMP expression and suppressing HSC activation in MASH.

## Discussion

We elucidate for the first time that RvD1 and its receptor FPR2 are decreased in human MASH livers. Moreover, we are the first to perform single-cell RNA sequencing in RvD1-treated MASH livers to confirm the beneficial effects of RvD1 on MASH. Mechanistically, we identify novel roles of RvD1 in Stat1/Cxcl10-mediated inflammation, ROS-mediated cell death, and MMP-mediated fibrosis regression in MASH.

Although it has been established that endogenous production of SPMs, including RvD1, become impaired in chronic diseases, including in MASH,^6,14,42^ their underlying mechanisms are not well understood. Along this line, we find that liver RvD1 levels are dramatically reduced in both MASH patients and MASH mice. Our *in vitro* studies with macrophage polarization provide new insight into the defective production of RvD1 in MASH livers. We show that pro-inflammatory M1-like macrophages have reduced capacity to produce RvD1, whereas anti-inflammatory M2-like macrophages generate more RvD-1 compared to unstimulated macrophages. Since M1-like macrophages are predominant in MASH, decreased RvD1 may be due to the abundance of pro-inflammatory macrophages with impaired capacity to biosynthesize RvD1. However, the mechanisms underlying varying levels of RvD1 production across different macrophage subsets need further clarification.

In addition to reduced RvD1, we also find that FPR2, one of the well-characterized RvD1 receptors, is decreased in MASH livers, indicating that RvD1-FPR2 signaling is suppressed in MASH. However, we do not observe a decrease in *Fpr2* mRNA in MASH livers (data not shown). Therefore, FPR2 may be regulated at a post-transcriptional level in response to MASH challenge. FPR2 is expressed in hepatocytes,^43^ myeloid cells (*i.e.*, macrophages, neutrophils),^44^ and T cells.^27^ FPR2 binds a wide range of ligands that trigger either a pro-inflammatory or an anti-inflammatory response.^43–47^ For instance, polypeptides, including serum amyloid A (SAA) and β-amyloid peptide 42 (Aβ_42_), elicit the pro-inflammatory effect of FPR2, while bioactive lipid molecules such as annexin A1 (ANXA1), lipoxin A4 (LXA_4_), and RvD1 trigger an anti-inflammatory signaling of FPR2.^46,47^ Accordingly, the role of FPR2 could be either beneficial or detrimental in MASLD/MASH. Lee et al.^43^ demonstrated that hepatocyte FPR2 suppressed cell death, inflammation, and fibrosis in CDAHFD-induced MASH mice, whereas Chen et al.^45^ showed that myeloid FPR2 accelerated macrophage infiltration into the liver and promoted inflammation by acting on SAA in HFD-fed mice. Given the complexity of FPR2 biology, the roles of FPR2 signaling in different cell types in the context of MASH warrant future investigation.

RvD1 can elicit beneficial effects in various liver injury models, including in MASLD, alcoholic hepatitis, and ischemia/reperfusion-induced injury.^17,48,49^ However, in those studies, single-cell RNA-seq was not performed to uncover the effect of RvD1 on immune and stromal cells. Our data are the first to apply single-cell RNA-seq to RvD1-treated FAT-MASH livers and demonstrate that RvD1 suppresses the activation of immune cells and HSCs. Suppressing the activation of nuclear factor-κB (NF-κB) was previously considered as an important mechanism by which RvD1 triggers anti-inflammatory activity and tissue repair in chronic inflammatory diseases, including in MCD-induced MASH.^17^ Here we demonstrate in the FAT-MASH model a new role of RvD1 in suppressing Stat1-mediated inflammatory response. Cxcl10, a Stat1-targeted inflammatory mediator, has been shown to promote liver inflammation in ischemia/reperfusion injury,^50^ enhance liver fibrosis in CCl_4_-injured livers,^51^ and exaggerate liver inflammation and fibrosis in FFC-induced MASH.^52^ Our study highlights the Stat1-Cxcl10 pathway as a novel mechanism by which RvD1 suppresses inflammation in MASH. In addition to suppressing inflammation, RvD1 treatment prevents apoptosis in our FAT-MASH mice. Hepatocyte apoptosis is an important feature of MASH and one of the primary causes of liver inflammation and fibrosis.^8^ Interestingly, our *in vitro* studies with primary hepatocytes show that RvD1 significantly prevents both palmitate-and Fas (Jo2)-induced apoptosis. It is notable that DEG analysis from single-cell RNA-seq reveals that RvD1 also suppresses the apoptotic process in immune cells, indicating that RvD1 has a broad anti-apoptotic activity in MASH.

In conclusion, using unbiased transcriptomic approaches, we establish that RvD1 administration alleviates MASH progression by suppressing inflammation and cell death, while enhancing fibrosis regression. Our study highlights the therapeutic potential of SPMs to boost tissue repair in MASH where other anti-inflammatory and repair pathways are insufficient.

## Abbreviations

MASLD: metabolic dysfunction-associated steatotic liver disease
MASH: metabolic dysfunction-associated steatohepatitis
SPM: specialized pro-resolving mediator
RvD1: resolvin D1
BMDM: bone marrow–derived macrophages
HSC: hepatic stellate cell
ALT: alanine aminotransferase
LPS: lipopolysaccharides

## Acknowledgments

We thank the Liver Tissue Cell Distribution System at the University of Minnesota for providing human normal and MASH liver tissues. We also thank Rima Weinberg for providing language help.

