## Supplementary Data for "Therapeutic Activity of Resolvin D1 (RvD1) in Murine MASH"

### Supplementary Materials and Methods

#### Identification and Quantification of Liver RvD1 by LC-MS/MS

Mouse and human liver tissue was subjected to targeted LC-MS/MS analysis essentially as described previously.[1] Tissue was placed in 1 mL in ice-cold methanol containing commercially available deuterium-labeled standards (d5-RvD2 and d8-15(S)-HETE; Cayman Chemical, Ann Arbor, MI), finely minced, then stored at -80°C. Immediately prior to solid phase extraction (SPE), samples were centrifuged (13,000 rpm; 10 min; 4°C). The supernatants were collected and promptly acidified prior to addition to methanol- and water-conditioned C18 SPE columns (Biotage, Uppsala, Sweden) as described in detail previously.[2] Lipid mediators were eluted from the columns with methyl formate which was then evaporated under $N_2$ gas. Samples were resuspended in methanol:water (50:50) prior to injection into the LC-MS/MS system. Lipid mediators were analyzed using a Shimadzu LC-20AD with a Kinetex Polar C18 HPLC column (100 x 3 mm; 2.6 µm) maintained at 59°C, coupled to a QTrap 5500 mass spectrometer (AB Sciex) operated in negative ionization mode. Samples were subjected to gradient elution with a constant 0.5 mL/min mobile phase delivery, initially consisting of 45:55:0.01 (v/v/v) methanol:water:acetic acid, which was ramped to 80:20:0.01 over 16.5 min, and then to 98:2:0.01 over 2 min. Data was acquired using Analyst (v.1.7.1) software.

Identification of RvD1 and 17-HDHA was performed using multiple reaction monitoring transitions and enhanced product ion mode. Mediators were identified by matching chromatographic peak retention time with that of synthetic standards run in parallel (+/- 0.1 min). Additional peak identification criteria included signal:noise ratio >5, at least 5 points across the baseline, and an on-column calculated concentration above the lower limit of quantification for each mediator. Matching of fragmentation spectra of samples to synthetic standards was also used. Quantification of identified compounds was performed by comparing samples to a 12-point standard curve of synthetic standards run in parallel and by calculating the extraction recovery from SPE using a blank sample (containing only methanol and internal standards) that was treated the same as the samples except for not undergoing SPE. Peak identification and quantification were accomplished using SCIEX OS (v.2.0.1).

#### Bulk RNA-Sequencing and Data Analysis

Mouse liver tissues were subjected to bulk RNA-seq analysis. Briefly, total RNA was extracted using the RNeasy Mini Kit (Qiagen, #74106). RNA-seq library construction was performed at Innomics Inc. with a standard polyA-enrichment protocol. The sequencing was performed on an Illumina HiSeq 4000 sequencer, and 150-bp paired-end reads were obtained.

For bulk RNA-seq data processing, reads were aligned to the mouse genome mm10 using STAR (v2.7.6a) with the default settings. Transcript assembly and differential expression analyses were performed using cufflinks (v2.2.1). Assembly of novel transcripts was not allowed (-G). Other parameters of cufflinks were the default setting. The summed FPKM (fragments per kilobase per million mapped reads) of transcripts sharing each gene_id was calculated and exported by the cuffdiff program. Differentially expressed genes (DEGs) were based on FPKM+1 values and determined by two-sided T-test P-value<0.05 and fold-change>1.25. Volcano plots for gene expression by fold change versus P-value were generated using R. Principal component analysis (PCA) was performed with Cluster 3.0 software. PC values were visualized with the plot3d function in the rgl package using R (v4.3.2) scripts. Gene set enrichment analysis (GSEA) (v4.2.3) was used to determine the statistically enriched gene sets by comparing the MASH and MASH+RvD1 groups. The curated C5: ontology gene sets were downloaded from https://www.gsea-msigdb.org/gsea/. The GSEA enrichment plot, normalized enrichment score (NES), and Q-value (FDR) were indicated for each enrichment test.

**Single-cell RNA-sequencing (scRNA-seq) and Data Analysis**

Mouse liver non-parenchymal cells (NPC) for scRNA-seq were isolated as previously described.[3] Briefly, the liver was perfused in situ with an HBSS buffer (without calcium and magnesium) containing 0.2 mg/mL EDTA, followed by sequential perfusion with an HBSS buffer (with calcium and magnesium) containing 0.4 mg/mL pronase (Sigma, P5147) and 0.2% collagenase type II (Worthington, LS004196). The liver was minced and further digested with an HBSS buffer (with calcium and magnesium) containing 0.2% collagenase type II, 0.4 mg/mL pronase and 0.1mg/mL DNase I (Roche, R104159001) in a 37$^{o}$C incubator with shaking for 30 min. Digestion was then stopped with DMEM containing 10% serum. The resulting liver cell suspension was passed through a 100 mm nylon cell strainer and centrifuged at 50 g for 3 min three times to remove hepatocytes. This NPC suspension was centrifuged at 800 g for 5 min. The NPC cell pellet was subjected to a red blood cell lysis buffer for 2 min. Upon the removal of red blood cells, the NPC cells were resuspended in HBSS, and subjected to density gradient centrifugation using 20% Optiprep (Axis Shield, #1114542) to remove dead cells. Cell viability was confirmed by trypan blue exclusion. scRNA-seq libraries were prepared using a Chromium Single Cell 3' Reagent kit (10x Genomics) according to the manufacturer's instructions. The libraries were sequenced on an Illumina NovaSeq 6000 sequencing system at the Mount Sinai Genomics Core Facility. The 10x Genomics Cell Ranger (v5.0.1) pipeline was used to process the data with the reference mm10.

The scRNA-seq filtered raw counts matrices were analyzed in Seurat (v5.0.1) using R (v4.3.2). During quality-control filtering, cells with less than 1,500 genes and the percentage of mitochondrial mRNAs more than 20% were removed. The filtered data were normalized using the NormalizeData function, then 2,000 of the most variable genes in each dataset were identified using the FindVariableFeatures function with the vst method. Different datasets were anchored with the FindIntegrationAnchors and IntegrateData functions and normalized with the ScaleData function. PCA was performed using the RunPCA function on the scaled expression data. Top informative PCs were selected (n=30) using the ElbowPlot function. The FindNeighbours, FindClusters, and RunUMAP functions were used to identify neighbors of cells, cluster cells, and UMAP (Uniform Manifold Approximation and Projection) virtualization. If not specified, the other settings followed the default values in Seurat. We tested resolution values from 0.5 to 1.0 in the FindClusters function and determined 0.7 as the best resolution value. Cells were annotated based on marker genes identified from the FindMarkers function and the known lineage markers from the literature. The quality of clusters was assessed by log(nCount_RNA) values. Clusters with low nCount_RNA values and expression of multiple lineage markers (likely due to contamination of ambient RNAs) were removed. A UMAP plot, feature plot, dot plot, and violin plot with cell annotations were virtualized using Seurat.

In each annotated cell type, the DEGs between MASH and MASH+RvD1 groups were determined by $P<0.05$ and percent of expression >0.25 in both groups. Gene ontology (GO) analysis was performed on these DEGs using the DAVID tool (https://david.ncifcrf.gov/tools.jsp), with all *Mus musculus* genes as a reference list.

CellPhoneDB[4,5] was performed for cell-cell interaction analysis using a curated database of ligand-receptor interactions (v5). Mouse genes were replaced by uniquely matched homologous human genes from the database at www.informatics.jax.org/downloads/reports/ HOM_MouseHumanSequence.rpt, and the cluster metadata file was created from the cell type annotations. CellPhoneDB analysis was performed using the cpdb_statistical_analysis_method (method 2) function on Python (v3.11.5) and Jupyter Notebook (v1.0.0). Other settings followed the default values. The output cell-cell interaction matrices were virtualized as heatmaps using the R packages pheatmap (v1.0.12) and ktplots (v2.0.0).

**Histological Analysis of FAT-MASH Mouse Livers**

Inflammation and fibrosis in FAT-MASH mouse liver tissues were analyzed by hematoxylin/eosin (H&E) and picro-Sirius Red staining, respectively. Inflammatory cells in H&E-stained liver section images were quantified as the number of mononuclear cells per field. Lipid droplet area in H&E-stained sections was quantified as % lipid droplet area of total area by ImageJ. For liver fibrosis evaluation, rehydrated liver sections were immersed in a solution containing saturated picric acid and 0.1% Sirius red (Sigma, Direct Red-80) for 1 h followed by counterstaining with 0.01% Fast Green for 1 h. Liver fibrosis was quantified as % Sirius red+ area of total area using Image J. For all the imaging analyses, five to ten fields per section per mouse were randomly chosen and quantified using the same ImageJ threshold settings.

**Mouse Plasma Analysis**

Mouse plasma levels of alanine aminotransferase (ALT) and cholesterol were measured using commercial kits (#A526-120 from TECO Diagnostics and #999-02601 from Wako Diagnostics, respectively).

**Preparation of Mouse Bone Marrow–Derived Macrophages (BMDMs)**

Bone marrow–derived macrophages (BMDMs) were isolated from the hind legs of 7-to-10-week-old C57BL/6J male mice as previously described.[1,6] In brief, the mouse was euthanized, and the skin was cut open to expose the hind legs. Cuts were then made through the hip and ankle joints to remove each leg. In the tissue culture hood, bone marrow was flushed out using a syringe and a 25-gauge needle containing BMDM media. Bone marrow cell suspension was filtered and cultured for 7–10 days in DMEM supplemented with 10% (vol/vol) FBS, 1% pen-strep, and 20% (vol/vol) L-929 fibroblast-conditioned media.

**Isolation of Primary Hepatocytes**

Primary hepatocytes were isolated from 7-to-10-week-old C57BL/6J mice as described previously.[7] Briefly, the mice were euthanized with isoflurane and a cannula was inserted into the vena cava to perfuse the liver with Hanks' Balanced Salt solution. Next, the portal vein was cut, and the liver was further perfused and digested with collagenase D (Roche, #11088858001). After the digestion, the livers were excised and minced in a petri dish containing DMEM. The digested liver tissue was passed through a 100 mm filter and centrifuged at 50 x g for 2 min. The cell pellets were resuspended in Percoll solution (Cytiva, GE17-0891-01) and then centrifuged at 200 x g for 10 min at 4°C. Supernatant was discarded and pelleted hepatocytes were washed once with DMEM. The hepatocytes were cultured in DMEM/F12 containing 10% (vol/vol) heat-inactivated FBS and 1% penicillin/streptomycin in collagen-coated plates, prepared with collagen coating solution (Sigma, C3867).

**Isolation of Kupffer Cells and Liver T Cells**

Kupffer cells and T cells were isolated from 7-to-10-week-old C57BL/6J mice as described previously.[6] Briefly, the mice were euthanized with isoflurane and a cannula was inserted into the inferior vena cava. After cutting the portal vein, the liver was perfused through the vena cava cannula with calcium- and magnesium-free Hanks' Balanced Salt solution (HBSS) buffer and then HBSS buffer (with calcium and magnesium) containing collagenase D (3.7 U/mouse, Roche). A tissue homogenizer was used to disrupt the digested liver, and the resulting cell suspension was centrifuged at 60 x g for 1 min to pellet hepatocytes. Kupffer cells and T cells contained in the supernatant fraction were then isolated by using anti-F4/80 (MACS, #130-110-443) and anti-CD90.2 (MACS, #130-121-278) microbeads, respectively, and cultured in RPMI media containing 10% (vol/vol) FBS and 1% pen-strep.

**Cell Culture and Treatments**

The experiments were conducted utilizing several murine primary cell types, including BMDMs, Kupffer cells, T cells, and hepatocytes. These cell types were isolated and cultured as described above. Additionally, LX2 cells, a human immortalized HSC cell line,[8] were cultured to validate some single-cell RNA-sequencing findings. The following treatments were used at the final concentration and times as indicated in the individual experiments in the results section: resolvin D1, RvD1 (Cayman Chemical, #10012554); recombinant murine Interferon gamma, IFNγ (Peprotech, #315-05); palmitic acid, PA (Sigma, P9767); recombinant transforming growth factor beta 1, TGFβ1 (R&D Systems, #240-B); lipopolysaccharide, LPS (*Escherichia coli* 0111:B4; Millipore-Sigma, L2630); recombinant tumor necrosis factor alpha, TNFα (R&D Systems, #410-MT); and anti-Fas antibody (Jo2) (BD Biosciences, #554254).

**Protein Extraction and Western Blot**

Whole protein lysates from cultured cells were extracted with 5% β-mercaptoethanol-containing 2X Laemmli buffer and then heated at 100°C for 7 min. Liver protein lysates were obtained using a RIPA lysis buffer. Protein concentration was determined using bicinchoninic acid (BCA) assay (Thermo Fisher Scientific, #23227). Briefly, 15–50 µg of protein was loaded onto 4–20% Tris-glycine gels (Invitrogen, XP04205BOX), electrophoresed, and transferred onto 0.45 µm nitrocellulose membranes (Bio-Rad, #1620115) for blotting. The membrane was blocked in non-fat dried milk 0.1% Tween20 Tris buffered saline (TBST) blocking solution, incubated with primary antibodies in Intercept blocking buffer, rinsed with TBST, and incubated with HRP conjugated secondary antibodies (Supplementary Table S2). After washing, the membrane was exposed to SuperSignal™ West Pico PLUS Chemiluminescent Substrate (Thermo Fisher Scientific, #34580) and developed.

**Quantitative RT-PCR**

Total RNA was extracted from cultured cells or mouse liver tissue using the RNeasy Mini Kit (Qiagen, #74106). Reverse transcription was performed with 0.5–1 µg RNA using the High-Capacity cDNA Reverse Transcription kit (Applied Biosystems, #4374967). Quantitative RT-PCR was performed in a volume of 10 µL using iQ SYBR Green Supermix (Bio-Rad, #1708886) and the Light Cycler 480 II Real-Time PCR system (Roche). Primer sequences are shown in Supplementary Table S3.

**Immunofluorescence (IF)**

Paraffin-embedded mouse liver tissue sections were deparaffinized in xylene and rehydrated in a graded series of ethanol. Sections were then antigen unmasked with a citrate-based solution (pH 6.0, Vector Laboratories, H-3300) in a pressure cooker for 10 min. After rinsing in PBS (pH 7.4) and blocking in 5% donkey serum, the sections were incubated with primary antibodies at 4°C overnight, followed by room-temperature incubation with secondary antibodies for 1 h (Table S2). Slides were then mounted with DAPI-containing mounting solution (SouthernBiotech, #0100-20). Three to five imaging fields were randomly obtained for each mouse liver sample using Zeiss fluorescence microscopy. In ImageJ, for each field of image, a threshold was set to reduce image background and the $MAC2^{+}DAPI^{+}$ areas were selected and analyzed for the mean fluorescence intensity (MFI) of p-Stat1 staining within them. The MFI from the three to five fields was averaged for each mouse. The same protocol was followed to analyze the MFI of ROS staining within $HNF4\alpha^{+}$ cells in mouse liver tissue.

**TUNEL Assay**

Liver sections were stained with TUNEL using the *In Situ* Cell Death Detection Kit, TMR red (Roche, #12156792910). Five images per mouse were captured using Zeiss fluorescence microscopy. Data was plotted as the average of the number of TUNEL+ cells per liver section for each mouse.

**Macrophage Polarization and RvD1 ELISA**

BMDMs were plated in a 24-well plate and polarized in M1 and M2. Briefly, BMDMs were polarized to M1 macrophages by incubation with 25 ng/mL of LPS (Sigma, #8630) and 20 ng/mL of murine IFNγ (Peprotech, #315-05) for 22 h. Macrophage M2 polarization was obtained by incubating BMDMs with 20 ng/mL of murine IL 4 (Peprotech, #214-14) for 22 h. Untreated macrophages were referred to as M0. Once polarized, macrophages were incubated with 10 µM docosahexaenoic acid (DHA) (Cayman Chemicals, #90310) in DMEM containing $Ca^{2+}$ for 1–2 h at 37°C. Supernatants were collected and assayed with the RvD1 ELISA kit (Cayman Chemical, #500380). The plate was read at a wavelength of 420 nm and data was calculated and plotted as pg/ml.

**Quantification and Statistical Analysis**

All results are presented as mean ±SEM. Statistical significance was determined using GraphPad Prism software. P values were calculated using the Student's t test for normally distributed data and the Mann-Whitney rank sum test for non-normally distributed data. One-way ANOVA with Dunnett post-test was used to analyze multiple groups with only one variable tested. Two-way ANOVA with Dunnett post-test was used to analyze more than two groups with multiple variables tested.

#### Data and Code Availability

The RNA-seq data has been deposited at the Gene Expression Omnibus (GEO), accession codes: GSE232780, GSE263768, and GSE263770. The deposited data will be publicly available as of the date of publication. This paper did not report original code. Any additional information required to reanalyze the data reported in this paper is available from the lead contact upon request.

### Supplementary Figure 1

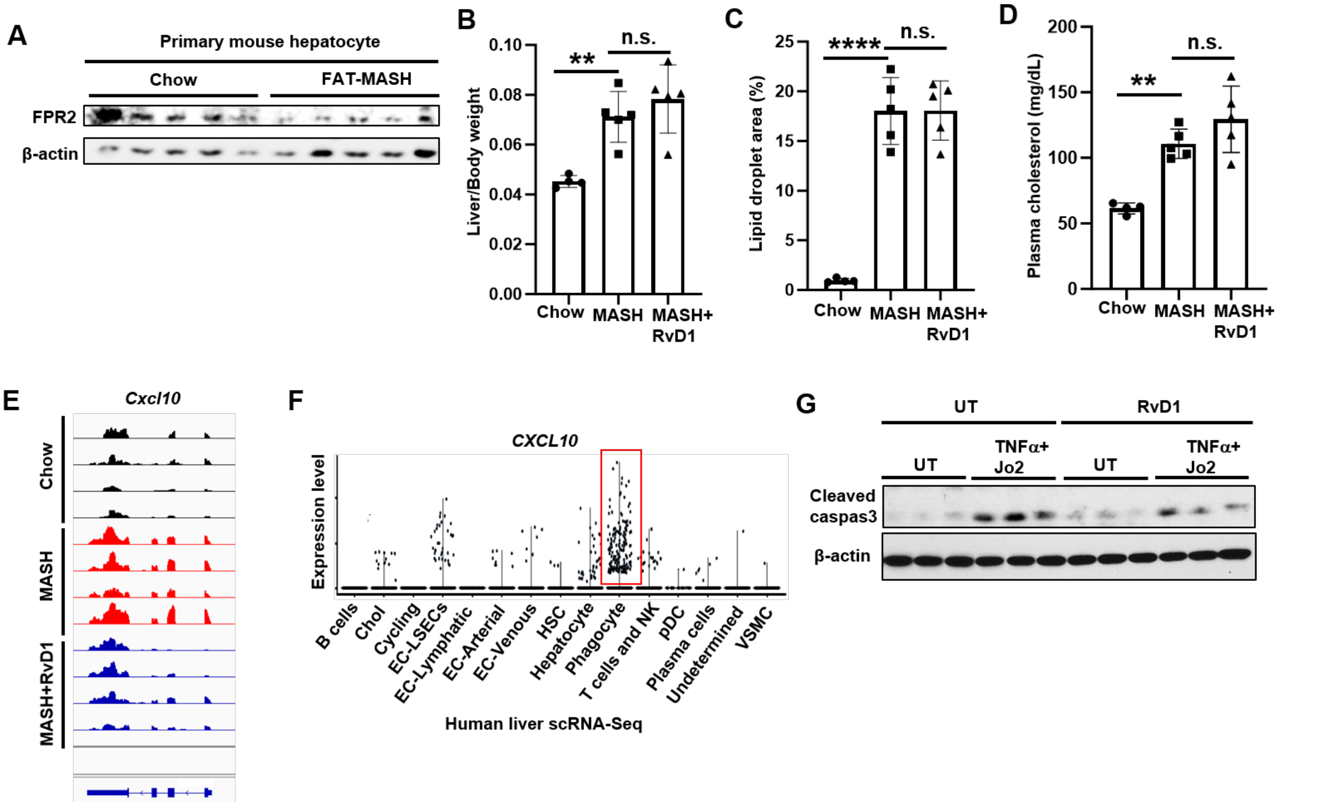

**Fig. S1. Effects of RvD1 on MASH livers and primary hepatocytes.** (A) Immunoblot of FPR2 in primary mouse hepatocytes isolated from chow- and FAT-MASH-fed mice (n=5 mice/group). (B) Liver/body weight ratios. (C) Percentage of lipid droplet area on HE-stained liver sections. (D) Plasma cholesterol. (E) RNA-seq track showing *Cxcl10* expression in mouse livers. (F) scRNA-seq on human livers[28] showing *CXCL10* expression in liver cell populations. Red box indicates phagocytes expressing higher levels of *CXCL10*. (G) Immunoblot of cleaved caspase-3 in primary hepatocytes untreated (UT) or treated with 50 ng/mL TNFα + 50 ng/mL Jo2 in the absence or presence of 50 nM RvD1 for 24 h. In B–D, n=4–5 mice/group; one-way ANOVA with Dunnett comparison **$p$<0.01, ****$p$<0.0001, n.s., no significance. Graph bars are presented as the mean ± SEM.

### Supplementary Figure 2

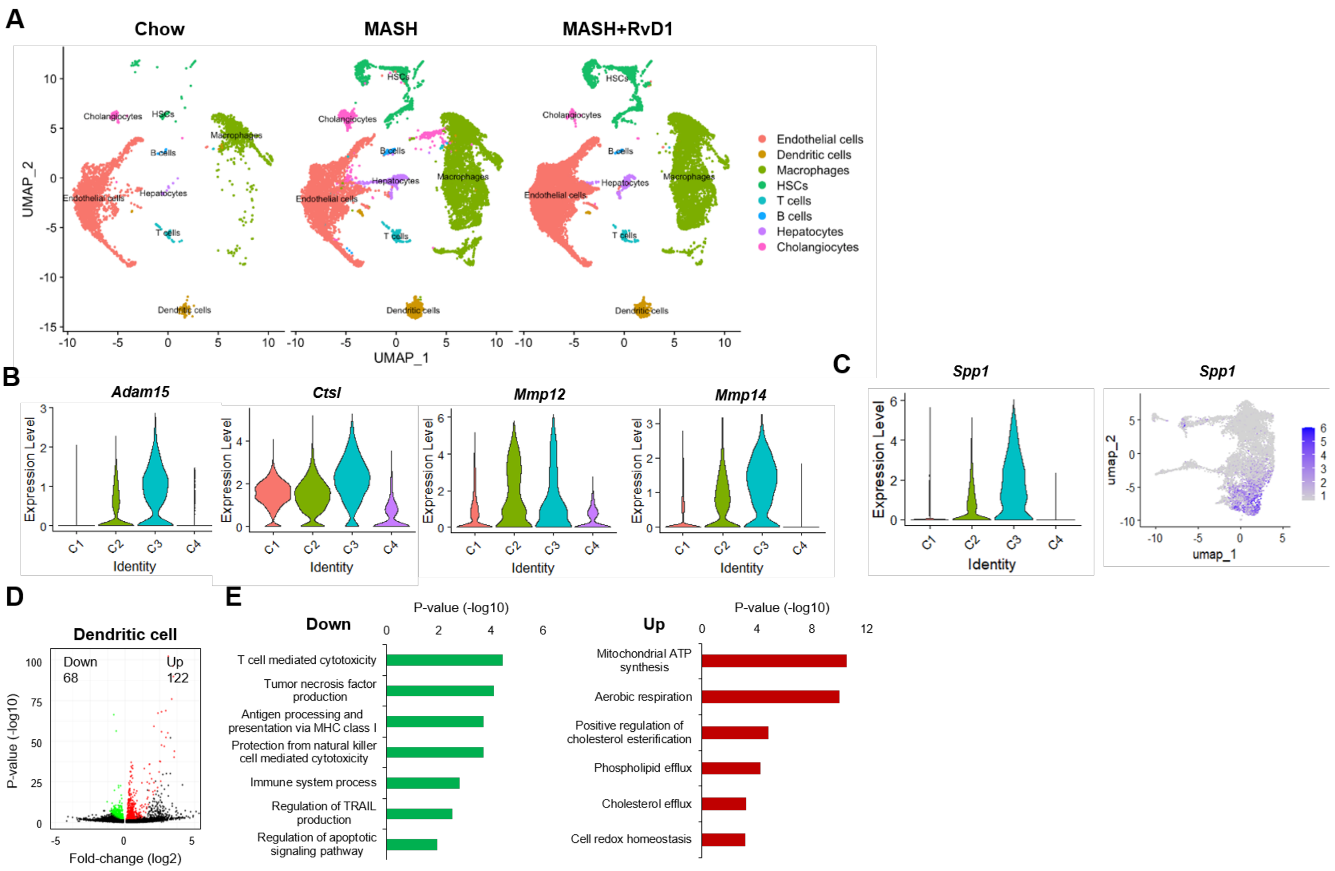

**Fig. S2. Single-cell RNA-seq analysis of liver cells in MASH mice.** (A) UMAP plot showing all cells split by groups of mice. (B) Feature plots showing expression of genes involved in collagen catabolic process. (C) Violin (left) and feature (right) plots showing *Spp1* expression. (D) Volcano plot showing DEGs in dendritic cells. (E) GO analysis for the downregulated (left) and upregulated (right) DEGs in dendritic cells.

### Supplementary Tables

#### Table S1. Antibodies used for immunoblot and Immunofluorescence.

| Antibodies | Supplier / Catalog. No |
|---|---|
| Rabbit anti-FPR2 (M-73) | Santa Cruz Biotechnology / #sc-66901 |
| Rabbit anti-iNOS | Abcam / #ab213987 |
| Rabbit anti-Arg1 | Cell Signaling Technology / #93668 |
| Rat anti-F4/80 | Biolegend / #123113 |
| Rabbit anti-phospho-STAT1 (Tyr701) | Cell Signaling Technology / #9167 |
| Rabbit anti-STAT1 | Cell Signaling Technology / #14994 |
| Rabbit anti-cleaved CASPASE-3 (Asp175) | Cell Signaling Technology / #9661 |
| Mouse anti-CHOP 3 (B-3) | Santa Cruz Biotechnology / #sc-7351 |
| Rabbit anti-beta-ACTIN (HRP) | Cell Signaling Technology / #5125 |
| Rat anti-MAC-2 | Cedarlane / #CL8942AP |
| Rabbit anti-HNF4a | Cell Signaling Technology / #3113 |
| Donkey anti-rabbit (AF647) | Invitrogen / #A31573 |
| Donkey anti-mouse (AF555) | Invitrogen / #A31570 |

#### Table S2. Primers used for qPCR.

| Organism | Gene | Forward (5-->3) | Reverse (5-->3) |
|---|---|---|---|
| Mouse | *Tnfa* | CTTCTGTCTACTGAACTTCGGG | CAGGCTTGTCACTCGAATTTTG |
| Mouse | *Ccl2* | TTAAAAACCTGGATCGGAACCAA | GCATTAGCTTCAGATTTACGGGT |
| Mouse | *Ifng* | ATGAACGCTACACACTGCATC | CCATCCTTTTGCCAGTTCCTC |
| Mouse | *Cxcl10* | CCAAGTGCTGCCGTCATTTTC | GGCTCGCAGGGATGATTTCAA |
| Mouse | *Acta2* | ATGCTCCCAGGGCTGTTTTCCCAT | GTGGTGCCAGATCTTTTCCATGTCG |
| Mouse | *Mmp2* | GAGATCTTCTTCTTCAAGGAC | AATAGACCCAGTACTCATTCC |
| Mouse | *Ddit3* | CCACCACACCTGAAAGCAGAA | AGGTGAAAGGCAGGGACTCA |
| Mouse | *Mmp9* | CTGGACAGCCAGACACTAAAG | CTCGCGGCAAGTCTTCAGAG |
| Mouse | *Mmp12* | CTGCTCCCATGAATGACAGTG | AGTTGCTTCTAGCCCAAAGAAC |
| Mouse | *Hprt* | TCAGTCAACGGGGGACATAAA | GGGGCTGTACTGCTTAACCAG |
| Human | *HPRT* | CCTGGCGTCGTGATTAGTGAT | AGACGTTCAGTCCTGTCCATAA |
| Human | *MMP2* | CCCACTGCGGTTTTCTCGAAT | CAAAGGGGTATCCATCGCCAT |

*Tnfa*, tumor necrosis factor-alpha; *Ccl2,* C-C motif chemokine ligand 2; *Ifng,* interferon gamma; *Cxcl10,* C-X-C motif chemokine ligand 10; *Acta2*, a-smooth muscle actin; *Mmp2, 9, 12,* matrix metallopeptidase 2, 9, 12; *Ddit3,* DNA damage inducible transcript 3; *Hprt*, hypoxanthine guanine phosphoribosyl transferase.

#### Table S3. Pathology of human MASH liver samples.

| ID | Age | Sex | Pathological diagnoses |
|---|---|---|---|
| 1 | N/A | N/A | Normal liver |
| 2 | N/A | N/A | Normal liver |
| 3 | N/A | N/A | Normal liver |
| 4 | N/A | N/A | Normal liver |
| 5 | N/A | N/A | Normal liver |

| | | | |
|---|---|---|---|
| 6 | N/A | N/A | Normal liver |
| 7 | N/A | N/A | Normal liver |
| 8 | N/A | N/A | Normal liver |
| 9 | N/A | N/A | Normal liver |
| 10 | N/A | N/A | Normal liver |
| 11 | 53 | F | MASH with fibrosis |
| 12 | 49 | M | MASH with fibrosis |
| 13 | 27 | F | MASH with fibrosis |
| 14 | 64 | F | MASH with fibrosis |
| 15 | 31 | M | MASH with fibrosis |
| 16 | 51 | M | MASH with fibrosis |
| 17 | 69 | M | MASH with fibrosis |
| 18 | 54 | M | MASH with fibrosis |
| 19 | 61 | M | MASH with fibrosis |
